# Niche mediated integrin signaling supports steady state hematopoiesis in spleen

**DOI:** 10.1101/2020.08.11.236083

**Authors:** Shubham Haribhau Mehatre, Irene Mariam Roy, Atreyi Biswas, Devila Prit, Sarah Schouteden, Joerg Huelsken, Catherine M. Verfaillie, Satish Khurana

## Abstract

Outside-in integrin signaling regulates cell fate decisions in a variety of cell types, including hematopoietic stem cells (HSCs). Our earlier published studies showed that interruption of Periostin (POSTN) and Integrin-**α**v (ITGAV) interaction induces faster proliferation in HSCs with developmental stage dependent functional effects. Here, we examined the role of POSTN-ITGAV axis in lympho-hematopoietic activity in spleen that hosts rare population of HSCs, the functional regulation of which is not clearly known. *Vav-iCre* mediated deletion of *Itgav* in hematopoietic system led to higher proliferation rates, resulting in increased frequency of primitive HSCs in adult spleen. However, in vitro CFU-C assays demonstrated a poorer differentiation potential following *Itgav* deletion. This also led to a decrease in the white pulp area with a significant decline in the B-cell numbers. Systemic deletion of its ligand, POSTN, phenocopied the effects noted in *Vav-Itgav*^*−/−*^ mice. Histological examination of *Postn* deficient spleen also showed increase in the spleen trabecular areas. Surprisingly, these were the myofibroblasts of the trabecular and capsular areas that expressed high levels of POSTN within the spleen tissue. In addition, vascular smooth muscle cells also expressed POSTN. Through CFU-S_12_ assays, we showed that hematopoietic support potential of stroma in *Postn* deficient splenic hematopoietic niche was defective. Overall, we demonstrate that POSTN-ITGAV interaction plays important role in spleen lympho-hematopoiesis.

## Introduction

In adult mammals, blood cell production takes place in the bone marrow (BM), wherein the hematopoietic stem cells (HSCs) reside and function, taking cues from extrinsic factors (1). During the onset of postnatal hematopoiesis, thymus takes up specialized function of lymphopoiesis, while spleen shows compartmentalisation into lymphopoietic and non-lymphopoietic zones (2). Both of these zones show developmental as well as structural distinctions and support remarkable differences in function. Although spleen acts as a secondary lymphoid organ that plays crucial role in both innate and adaptive immune system functions, neonatal splenectomy did not alter essential hematopoietic processes (3). Apart from the active hematopoietic events during embryonic development or immune-activation, spleen undertakes extramedullary hematopoiesis (EMH) during BM failure in several pathological conditions, pregnancy and blood loss (4).

It is believed that EMH involves migration of HSCs from their normal niches; and homing, proliferation and differentiation supported by the extramedullary sites (5). In this process, a role of activated macrophages in modulating the capacity of spleen niche to support incoming HSCs was demonstrated. Silencing of the receptor for macrophage-colony stomulating factor (M-CSF) through nanoparticle encapsulated siRNA that caused decrease in the macrophage numbers, led to a reduction in spleen resident HSCs (6). This study further showed that M-CSF silencing selectively decreased the expression of vascular cell adhesion molecules 1 (VCAM-1), primarily expressed by macrophages in spleen. VCAM-1 is a ligand for integrin heterodimer VLA-4 (Very Late Antigen-4, integrin-**α**4**β**1) (7), well known for its role in homing, maintenance and functioning of HSCs in the BM (8). Inhibition of VLA-4 using small molecule inhibitors (9) or specific antibodies (10) induced mobilization of HSCs. Integrins are one of the most important classes of cell surface receptors that mediate HSC interaction with their niche through outside-in and inside-out signaling mechanisms (11, 12).

Although it was reported that HSCs might follow distinct mechanisms for homing and functioning in the BM and spleen niche (13), evidence suggested that integrin signaling was important for the establishment of hematopoietic activity in embryonic spleen (14). Even homing of transplanted HSCs into spleen, similar to the BM, was shown to be mediated via interaction of HSCs with their niche through integrins (8). Therefore, the importance of cell adhesion, mediated by inside-out integrin signaling, in hematopoietic processes in BM and spleen has been known. In contrast, the evidence of involvement of ligand mediated outside-in integrin signaling is not clearly elucidated. The importance of Osteopontin (OPN), a known factor that elicits outside-in integrin signaling as a negative regulator of HSC function, has been well established (15, 16). OPN binds to a variety of integrin heterodimers, most important being the ones with integrin-**α**v (ITGAV) as one of the monomers (17). In our recent findings, Periostin (POSTN) that shows binding-specificity to ITGAV-containing integrin heterodimers (18), is demonstrated as a negative regulator of HSC proliferation (12). However, another study reported the involvement of integrin-**α**v**β**3, synergising the effect of interferon-**γ** (IFN-**γ**) on HSC proliferation, resulting in the loss of hematopoietic function (19). Therefore, the involvement of ligand mediated outside-in integrin signaling in HSC function has not been well established yet.

Spleen is a major site for extra-medullary hematopoiesis (EMH), where HSCs mobilised from the BM play important role. In this context, a role of spleen resident HSCs, a rather rare population, is not well documented. It is important to note that at the cellular level, spleen resident HSCs have been reported to possess function, equivalent to the BM HSCs (20). However, their function in maintaining the supply of mature blood cells in not clearly known and regulatory pathways remain elusive. Here we show that POSTN, expressed majorly in the myofibroblasts of the splenic capsule and trabeculae, is involved in spleen lympho-hematopoietic activity. Lack of its receptor, ITGAV, led to increased proliferation rate in HSCs and decrease in the B-cell population with concomitant increase in the megakaryocyte numbers. POSTN deficiency largely mimicked the phenotype with additional defects in spleen hematopoietic niche that led to poorer support for the incoming hematopoietic progenitors. Overall, we show that ITGAV-mediated outside-in integrin signaling plays an important role in the steady state hematopoietic activity in spleen.

## Materials and methods

### Mice

Twelve to sixteen week-old FVB/NJ, C57BL/6J-CD45.2 (*Centre d’Elevage* R. *Janvier,* Le Genest-St Isle*, France),* and *Postn−/−* mice (21) were maintained in the animal facilities at IISER TVM and KU Leuven. *Itgav*^*flox/flox*^ mice (22) were crossed to *Vav-iCre* mice (23) from Thomas Graf, Centre for Genomic Regulation, Barcelona) to obtain *Vav-iCre;Itgavfl/fl* mice. Genotyping was performed on genomic DNA from tail tips, as described before (12). *Vav-iCre-;Itgav*^*fl/fl*^ and *Vav-iCre*^*+*^*Itgav*^*+/+*^ littermates were used as controls. During the experiments, mice were maintained in isolator cages, fed with autoclaved acidified water and irradiated food ad libitum. All experiments were approved by the Institutional Animal Ethics Committee of IISER Thiruvananthapuram and KU Leuven.

### Flow cytometry

Chimerism and lineage analysis of BM derived cells was performed by flow cytometry, using specific antibodies. Cell cycle analysis was performed by Hoechst 33342 (Ho) and Pyronin Y (PY) staining, in addition to labelling with HSPC (Lin-Sca-1^+^c-kit^+^ cells) or HSC (CD150^+^CD48-Lin-Sca-1^+^c-kit^+^) markers. Complete list of antibodies used for these experiments is provided in Table 1. Details of the procedures followed can be found in Supplementary methods section.

### Quantitave RT-PCR

Quantitative RT-PCR was performed using standard protocols. Details of the procedures, reagents and equipment used are provided in supplementary methods. List of primers used is provided in Table 2.

### Colony-forming unit-spleen assay

Recipient *Postn*^*+/+*^ and *Postn*^*−/−*^ mice were lethally (10Gy) irradiated and injected with 1×10^5^ control FVB/NJ mouse BM derived cells. Three recipient animals were used for each donor mouse and the experiment was repeated thrice. Recipients were sacrificed 12 days after injection. Spleen colonies were visualized and counted as described before (24).

### Histological and immunofluorescence studies

Spleen tissues fixed with 4% paraformaldehyde (PFA) were cut into 10**μ**m thick sections. These sections were used for hematoxylin and eosin (H&E) or immunofluorescence studies. Details of procedures and imaging tools used for analysis are provided in supplementary methods. For each experiment, appropriate matched-isotype antibody controls were used (shown in Suppl. Fig. 1).

**Figure 1.**
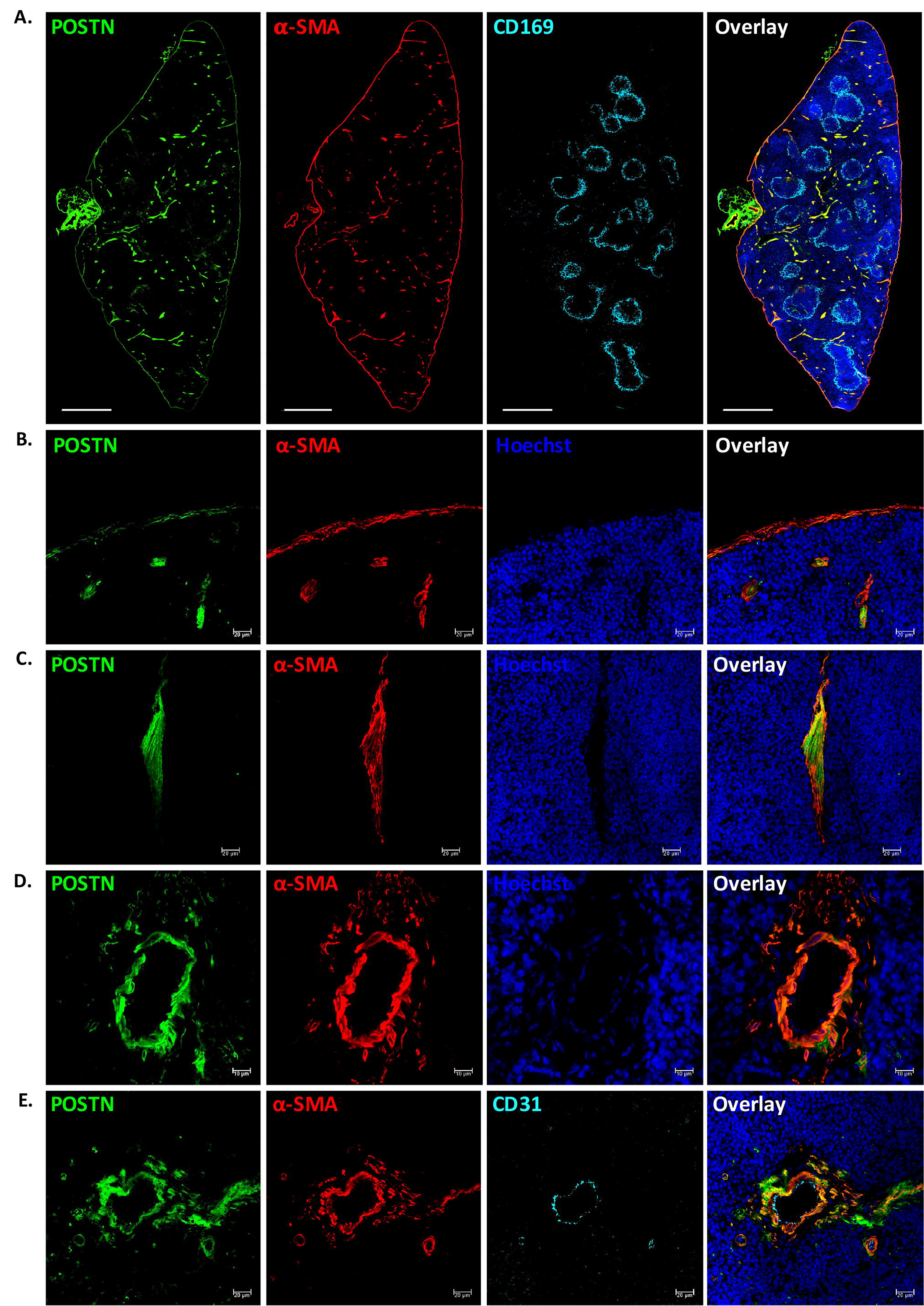
POSTN is expressed in myofibroblasts in adult spleen. (A) POSTN expression in spleen tissues was examined in the adult spleen tissues using immunohistochemistry on 10μm cryo-sections. WP areas were identified by using macrophage marker CD169; trabecular, capsular and vascular regions were identified by using antibodies against α-SMA. Specific antibodies were used to identify the cells expressing POSTN, nuclear counterstaining was done using Hoechst 33342. The sections were visualised and tile scanning was performed on confocal microscopes. (n=2, N=8, scale bar=0.5mm). (B-D) Regions with compelling levels of POSTN expression; namely, the capsule (B), Trabeculae (C), and vasculature (D) were then specifically analyzed by immuno-staining with myofibroblast/ smooth muscle cell marker α-SMA. (n=5, scale bar= 20μm for panel B,C and 10μm for panel D). (E) The endothelial and smooth muscle lining of the vessels was identified using endothelial marker CD31, in addition to smooth muscle cell marker α-SMA to identify the source of Postn (n=3, scale bar= 20μm).

### Statistical analysis

All data are represented as mean±SEM. Comparisons between samples from two groups were done using unpaired Student’s *t* test. For multiple comparisons, one-way ANOVA followed by Tukey Kramer post hoc test was used. Statistical analyses were performed with Microsoft Excel or GraphPad Prism 6. For all analyses, p-values ≤0.05 were accepted as statistically significant.

## Results

### POSTN is expressed by capsular and trabecular myofibroblasts

We first examined the spatial localisation of POSTN in the spleen tissue by performing immunohistochemistry using anti-POSTN antibodies. As spleen is functionally divided into two major regions, white pulp (WP; for lymphopoiesis) and red pulp (RP; for blood filtration), we tested if the expression of POSTN was restricted to one of these regions. We used anti-CD169 antibodies to detect macrophages of the marginal zone that clearly demarcate WP from RP (25, 26). We performed tile scans of the transverse sections of spleen and observed that the expression of POSTN was mostly limited to the RP (Suppl. Fig. 2). However, modest level of POSTN expression was observed in the vascular region of WP as well. Preliminary observations indicated that POSTN expression might be associated with the capsular, trabecular and vascular regions of the spleen tissues. To confirm this, we used anti-α-smooth muscle actin (α-SMA) antibodies to identify specific cells and splenic regions with POSTN expression (Fig. 1A). We confirmed that POSTN expression was mostly but not completely confined to the RP area. We also observed that the myofibroblastic cells of the capsular (Fig. 1B) and trabecular regions (Fig. 1C), as well as the smooth muscle cells of the vasculature (Fig. 1D) showed significant levels of POSTN expression. Although capsular myofibroblasts expressed lower levels of POSTN, all α-SMA+ cells in the spleen tissue were seen to express POSTN. Our recent findings showed that vascular endothelium was the key source of POSTN in the fetal liver (FL) HSC niche, with no significant expression seen in the vascular smooth muscle cells (27). Therefore, we tested if the vascular endothelium, in addition to the smooth muscle cells associated with vasculature expressed POSTN (Fig. 1E). We noted that the endothelial lining (CD31+) of the blood vessels did not express POSTN while the surrounding α-SMA+ cells showed high level of POSTN expression. These results show that cellular source of POSTN may vary between different tissues or developmental stages.

**Figure 2.**
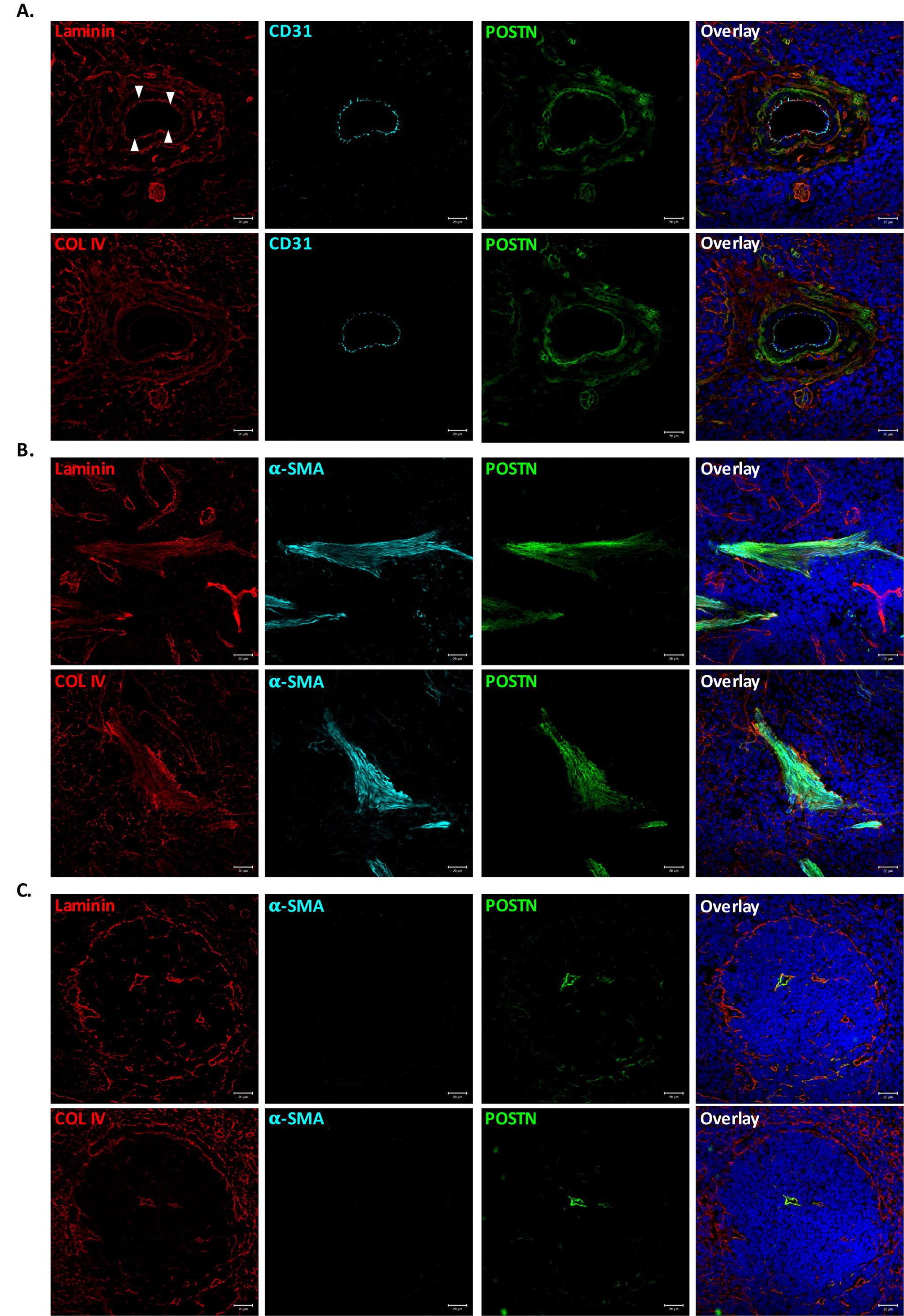
POSTN colocalizes with ECM proteins close to myofibroblasts. Immunostaining experiments were performed to examine co-localization of POSTN with ECM proteins laminin and Collagen IV in various anatomical locations of spleen tissue. (A) Vascular region of spleen identified by CD31 immunostaining. Colocalization of POSTN with laminin (upper panel) and Collagen IV (lower panel) was performed using specific antibodies. (B) Trabecular area is identified by α-SMA staining of myofibroblasts. Co-localization of POSTN along with laminin (upper panel) and Collagen IV (lower panel). (C) White pulp area, identified by the laminin (upper panel) and Collagen IV (lower panel) expression on the periphery and in red pulp. identification of smooth muscle cells/myofibroblasts and POSTN using specific antibodies. n=3, Scale bar=20**μ**m

As POSTN is an extracellular matrix (ECM) binding protein, we examined its association with two of the most common ECM proteins in spleen tissue, Laminin and Collagen IV, in different anatomical locations within the spleen tissue (Fig. 2). Distribution pattern of the two proteins in vascular region was similar, except for the vascular basal lamina that was composed mainly of laminin. Importantly, there was no clear preferential co-localization of POSTN with either laminin (Fig. 2A, upper panel) or COL IV (Fig. 2A, lower panel). Rather, it appeared to be determined by the closeness to the POSTN expressing cells. Therefore, ECM associated POSTN was more closer to the smooth muscle cells underneath the endothelial layer (white arrows) of the vessel. In the case of trabecular (Fig. 2B, upper and lower panel) as well as lymphopoietic WP area (Fig. 2C, upper and lower panel), POSTN localisation was not determined by the ECM proteins rather by the presence of smooth muscle cells/myofibroblasts in the region.

### ITGAV and ITGB3 are expressed in the spleen hematopoietic stem cells

We next performed flow cytometry analysis to examine the expression of ITGAV and ITGB3 in different sub-populations of spleen resident hematopoietic stem and progenitor cells (HSPCs; Figure 3). On the basis of expression of SLAM markers, CD150 and CD48, lin-c-kit^+^Sca-1^+^ (LSK) population was subdivided into four sub-populations; CD150+CD48-(P1), CD150+CD48+ (P2), CD150-CD48+ (P3) and CD150-CD48-(P4) (Fig. 3A). Subsequently, the expression of ITGAV (Fig. 3B, upper panel) and ITGB3 (Fig. 3B, lower panel), in each of these populations was examined. Results showed that among all the sub-populations, primitive HSCs (P1) expressed the highest level of ITGAV expression (55.54±6.26%). However, the short-term (ST-) HSCs (P2) as well as other multipotent progenitors (P3 and P4) also expressed significant levels of the protein (Fig. 3C). As ITGAV has been shown to partner with ITGB3 to form a heterodimeric integrin receptor in a variety of cell types (28), including a number of cancers (29), we investigated the expression of ITGB3 in the stem cell sub-populations. We observed that the proportion of most primitive HSCs (P1) that expressed ITGB3 was higher than both CD150+CD48+ and CD150-CD48+ LSK sub-populations (Fig. 3C). Thus, a significant proportion of all sub-populations, most importantly the primitive HSCs, expressed ITGAV and ITGB3. To confirm the co-expression of ITGAV and ITGB3 on primitive HSCs, we repeated flow cytometry experiments. The primitive HSCs (P1) was analyzed for the expression of both of the integrin chains (Fig. 3D,E). We observed that 36.98±3.63% of the most primitive spleen resident HSCs co-expressed ITGAV and ITGB3. Of note, 79.54±4.8% of the HSCs that expressed Itgav also expressed ITGB3 (Fig. 3E).

**Figure 3.**
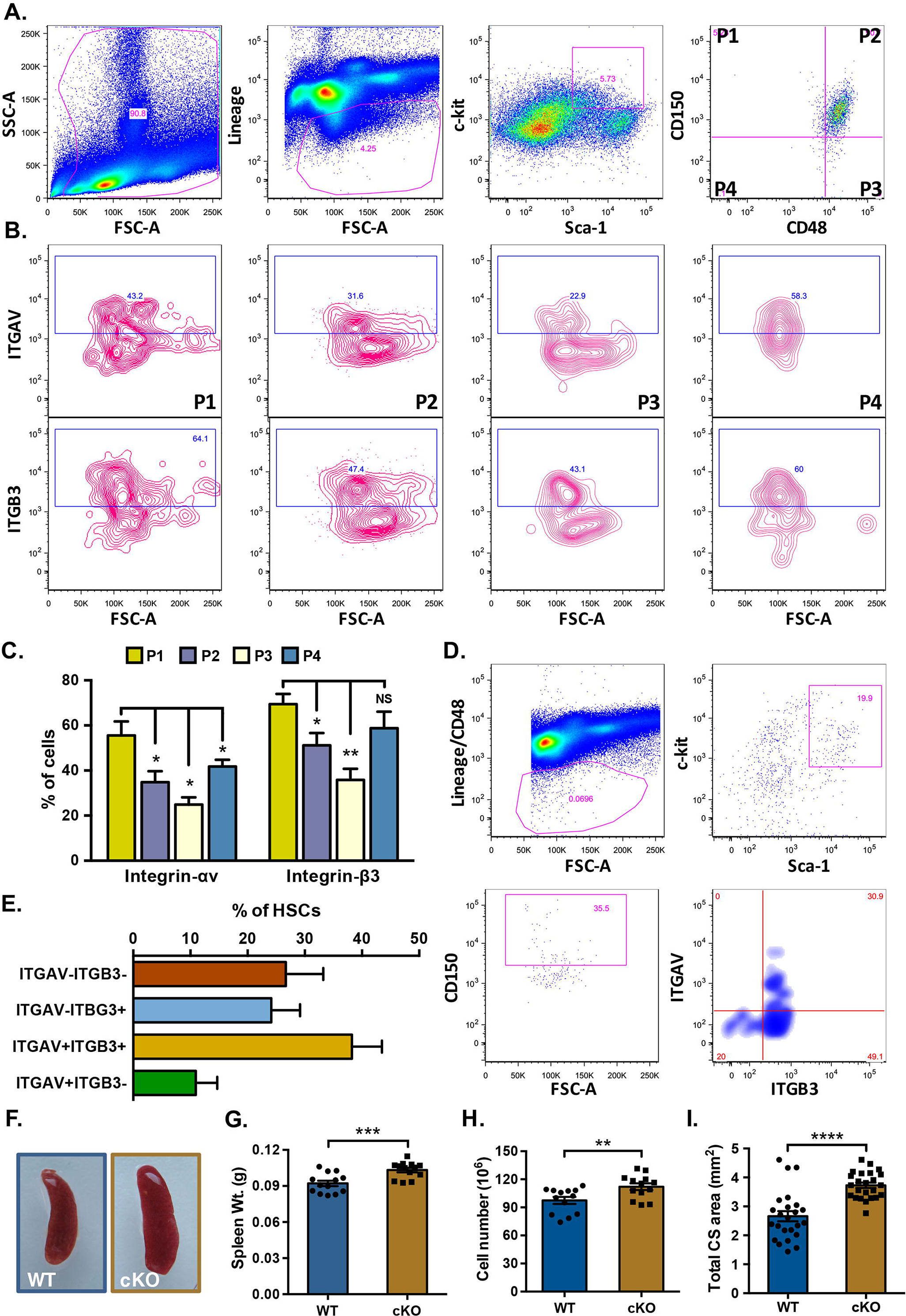
Expression of ITGAV and ITGB3 in hematopoietic stem cell sub-populations in spleen. Adult spleen derived MNCs were analyzed for the cell surface expression of ITGAV and ITGB3 on various HSC sub-populations by flow cytometry. (A) The LSK population from the spleen MNCs was further divided into four sub-populations on the basis of expression of SLAM markers CD150 and CD48. (B) All four sub-populations of the LSK cells were examined for the expression of ITGAV (upper panel) and ITGB3 (lower panel), separately. (C) Comparison of ITGAV and ITGB3 expression on different HSC sub-populations. Proportion of each LSK sub-population based on CD150 and CD48 that expressed ITGAV or ITGB3 was plomed. One-way ANOVA followed by Tukey-Kramer Post Hoc test was performed * p<0.05, ** p<0.01. (D) Co-expression analysis of ITGAV and ITGB3 on the primitive HSCs. The adult spleen derived HSCs were identified as lin-CD48-c-kit^+^Sca-1^+^CD150^+^ cells and were examined for the expression of ITGAV and ITGB3 based on flow cytometry. (E) Proportion of primitive HSCs expressing ITGAV and ITGB3 in combination or separately. (F) Gross morphology of the spleen from *Vav-Itgav+/+* (WT) and *Vav-Itgav*^*−/−*^ (conditional knockout; cKO) mice obtained following crossing *Vav-*iCre and *Itgav*^*fl/fl*^ lines. (G) Comparison of whole spleen weight from WT and cKO mice. (H) Comparison of total spleen cellularity (total mononuclear cells) in the WT and cKO mice. (I) Cross-sectional area quantified using transverse sections through the spleen tissues. Eight 10**μ**m thick sections (every sixth section used for analysis), were analyzed for total cross-sectional area. n=4-5, N=24, * p<0.05, ** p<0.01, *** p<0.001, **** p<0.0001, ns indicates not significant.

In order to understand if POSTN-ITGAV interaction played any role in spleen lympho-hematopoietic activity, we conditionally deleted *Itgav* in hematopoietic cells by crossing *Itgav*^*fl/fl*^ (22) with *Vav-*iCre mice (23). Genotyping was performed on the DNA isolated from tail tip tissues of adult animals and *Vav-Itgav+/+* and *Vav-Itgav*^*−/−*^ mice were identified and used further analysis (Suppl. Fig. 3A-C). Initial observations suggested little change in the gross morphology of spleen, upon *Itgav* deletion (Fig. 3F). However, we observed a modest increase in spleen weight (Fig. 3G) and total cellularity (Fig. 3H). Analysis of the transverse sections through the tissue showed an increase in the cross-sectional area (Fig. 3I).

### *Itgav* deletion leads to increased HSC proliferation and poorer lymphopoietic function

To examine the effect of loss of ITGAV in spleen hematopoietic system, we compared *Vav-Itgav+/+* and *Vav-Itgav*^*−/−*^ mice for the frequency of HSPCs (lin-c-kit+ cells), LSK cells and primitive HSCs in spleen tissues (Fig. 4A-C, Suppl. Fig. 4A). Although we did not observe any significant change in the frequency of lin-c-kit+ cells (Suppl. Fig. 2A), we did observe a significant increase in the frequency of LSK cells (Fig. 4B) as well as primitive HSCs (Fig. 4C) following *Itgav* deletion. We then examined if this increase in the frequencies of HSPC populations was accompanied by an altered proliferation status. Cell cycle analysis of the control spleen derived mononuclear cells (MNCs), using Hoechst and Pyronin Y staining (Fig. 4D,E), showed 72.01±6.74% of the spleen resident primitive HSCs to be quiescent (in G0 stage). Upon *Itgav* deletion, we observed a significant decrease in the proportion (49.64±2.37%) of quiescent HSCs (Fig. 4E). No significant difference upon *Itgav* deletion was observed in the proportion of spleen HSCs in G1 phase of cell cycle (18.25±5.4% in WT and 25.65±4.87% in *Vav-Itgav*^*−/−*^). However, we observed an increase in the proportion of cells in S phase, from 4.83±1.22% to 11.85±1.39% and in G2/M phase, from 4.91±0.65% to 12.85±1.9% upon *Itgav* deletion in hematopoietic system (Fig. 4E). These results indicate that the loss of ITGAV in the hematopoietic system leads to increased proliferation rates. We next examined a possible impact of increased proliferation rates on the function of stem cell population by performing methyl cellulose based colony assays (Fig. 4F,G). Results showed no difference in total colonies formed (Fig. 4F) but the number of CFU-GEMMs per spleen (Fig. 4G) was significantly decreased, showing a functional decline in more primitive progenitors in the spleen, following *Itgav* deletion. Our earlier published results showed significant loss of long-term engraqment potential in the *Vav-Itgav*^*−/−*^ HSCs from BM, with concomitant loss of lymphocytes in peripheral blood (12). Here, we performed experiments to understand if *Itgav* deletion led to any loss of lympho-hematopoietic activity in spleen. We first performed H&E staining based histological analysis on 10**μ**m thick spleen sections (Fig. 4H) and observed a significant decrease in the WP area (Suppl. Fig. 4B) and a corresponding increase in the RP area (Suppl. Fig. 4C) upon *Vav*-iCre induced *Itgav*-deletion. This resulted in an overall increase in the RP/WP ratio in the spleen tissues from *Vav-Itgav*^*−/−*^ mice (Fig. 4I). Splenic WP area is associated with lymphopoietic activity. To confirm any change in T- or B-lymphopoietic activity, we performed flow cytometry analyses of the MNCs from spleen tissue (Fig. 4J-M). We observed increase in the myeloid population (CD11b/Gr-1+) in *Vav-Itgav*^*−/−*^ spleen (Fig. 4K). While no difference was observed in T-cell population (CD4/CD8+; Fig. 4L), *Itgav*-deletion led to a significant decrease in the B-cell population (B220+; Fig. 4M). Careful observations on the spleen tissue sections from *Vav-Itgav*^*−/−*^ mice showed an increase in the megakaryocyte frequency compared to the control (Suppl. Fig. 4D,E).

**Figure 4.**
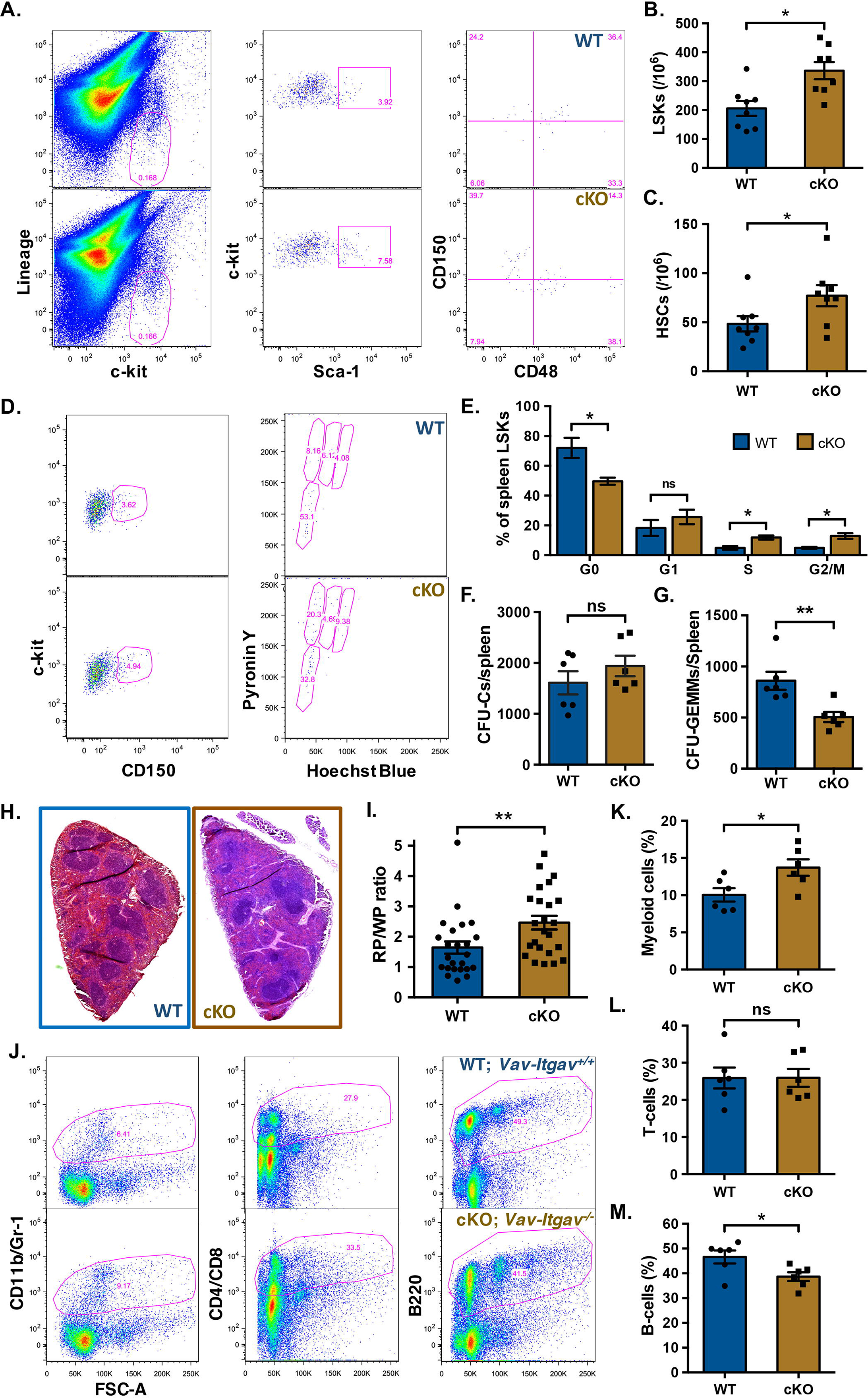
Increased frequency of phenotypic HSCs and poor B-lymphopoiesis in*Vav-Itgav*^*−/−*^ Spleen. (A-C) Mononuclear cells from *Vav-Itgav+/+* (WT), *Vav-Itgav*^*−/−*^ (cKO) spleen tissues were used for the analysis of HSC frequency by flow cytometry. Quantification of frequency of hematopoietic stem and progenitor cell populations in the spleen from WT and cKO mice; (A) lin-c-kit^+^ cells, (B) LSK cells, (C) primitive HSCs (n=8). (D,E) Cell cycle analysis of the primitive HSCs population from spleen by Hoechst 33342/pyronin Y staining. Comparison of the proportion of spleen LSK cells in various stages of cell cycle is shown (n=6). (F,G) CFU-C assay performed on total splenocytes to compare the frequency of hematopoietic progenitors. Total number of CFU-Cs (F) and CFU-GEMMs (G) were compared between WT and cKO mice (n=6). (H) Spleen tissues isolated from *Vav-Itgav*^*+/+*^ (WT) and *Vav-Itgav*^*−/−*^ (cKO) animals were harvested. The formalin-fixed, paraffin-embedded tissues were used to cut 10**μ**m sections that were used for H&E staining. (I) The ratio between the total cross-sectional area under red pulp (RP) and white pulp (WP) was compared between WT and cKO spleen tissues (n=4, N=24). (J-M) Mononuclear cells harvested from WT and cKO spleen tissues were used for flow cytometry analysis. The proportion of CD11b/Gr-1^+^ myeloid cells (K), CD4/CD8^+^ T-cells (L) and B220^+^ B-cells (M) was compared between WT and cKO mice derived spleen tissues (n=6, t test). Unpaired two tailed Student’s t-test was performed. * p<0.05, ns indicates not significant.

### *Postn* deletion partially mimics the effects observed in *Vav-Itgav*^−/−^ mice

To confirm that the effects of *Itgav* deletion reflected the importance of POSTN-ITGAV interaction specifically, we analysed spleen hematopoietic system following *Postn* deletion. Surprisingly, unlike in the case of *Vav-Itgav*^*−/−*^ mice, *Postn*^*−/−*^ mice showed significantly smaller spleens (Fig. 5A), as can be seen through a significant decrease in spleen weight (Fig. 5B), total cellularity (Fig. 5C) as well as average cross-sectional area (Fig. 5D, Suppl. Fig. 5A). Next, we examined the frequency of HSPCs (lin-c-kit+ cells), LSKs (lin-c-kit^+^Sca-1^+^ cells) and primitive HSCs (CD150^+^CD48-LSKs) in total spleen MNCs (Fig. 5E-G, Suppl. Fig. 5B). While we did not observe any change in the HSPC population (Suppl. Fig. 5B), there was a significant increase in the frequency of LSK cells (Fig. 5F) and primitive HSCs (Fig. 5G). We then asked if this increase in the stem cell frequency was linked to altered proliferation status (Fig. 5H,I). The LSK population was analyzed for cell cycle status using Hoechst 33342 and Pyronin Y staining (Fig. 5H, gated on LSK cells). Although we did not observe any significant change in the proportion of quiescent LSK cells (G0 stage), we observed a decrease in the cells in G1 (from 24.29±2.99% to 13.05±1.19%; Fig. 5I). Concomitantly, we found increase in the proportion of cells in S (from 9.15±0.83% to 14.04±1.38%) as well as G2/M phase (from 11.51±1.74% to 20.01±1.36%) of cell cycle. These results demonstrate faster proliferation of the spleen resident hematopoietic progenitors, as observed in the case of *Vav-Itgav*^*−/−*^ mice.

**Figure 5.**
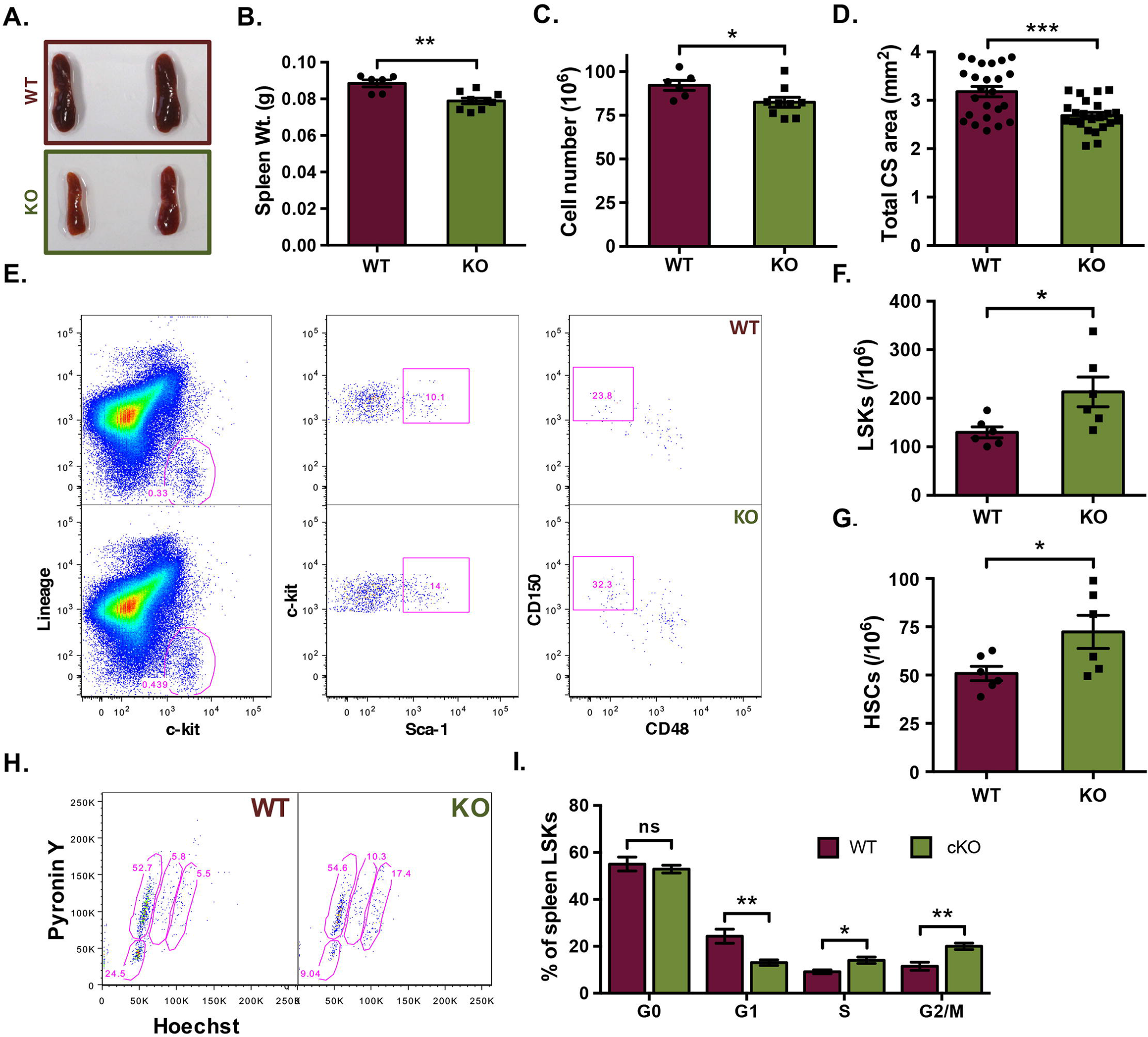
Loss of POSTN expression affects proliferation of hematopoietic progenitors in spleen. (A) Spleen tissues isolated from *Postn*^*+/+*^ (WT) and *Postn*^*−/−*^ (KO) mice were harvested. (B) Comparison of whole spleen weight from WT and KO mice. (C) The total number of cells from spleen from WT and KO mice. (D) Comparison between the cross-sectional area of WT and KO spleen. The formalin-fixed, paraffin-embedded tissues were used to cut 10**μ**m sections that were used for H&E staining and analysis. (E-G) Quantification of frequency of various hematopoietic stem and progenitor cell populations in the spleen from WT and KO mice; (F) LSK cells, (G) primitive HSCs. (H) Cell cycle analysis of the spleen LSK cells by Hoechst 33342/pyronin Y staining. Total MNCs from spleen tissues were used to first label the cells with lineage, Sca-1 and c-kit antibodies followed by Hoechst 33342/pyronin Y staining. Samples acquired on a flow cytometer were analysed and LSK cells were gated for further analysis for Hoechst 33342 and pyronin Y intensity. (I) Comparison of the proportion of spleen LSK cells in various stages of cell cycle (n=6). Unpaired two tailed Student’s t-test was performed. * p<0.05, ** p<0.01, ***p<0.001. ns indicates not significant.

Through careful histological analyses, we observed that *Postn* deletion led to increase in the trabecular area (Fig. 6A,B). In addition, as in the case of *Vav-Itgav*^*−/−*^ mice, we observed a significant increase in the RP area (Fig. 6C; Suppl. Fig. 6A), with a corresponding decrease in the WP area (Fig. 6C, Suppl. Fig. 6B) resulting in overall increase in the RP/WP ratio (Fig. 6D). However, *Postn* deletion did not result in any change in the proportion of myeloid (CD11b/Gr-1^+^; Fig. 6E), T-(CD4/CD8^+^; Fig. 6F), or B-cell (B220^+^; Fig. 6G) populations. From H&E stained sections of *Postn*^*−/−*^ spleen tissues, we observed a similar increase in the megakaryocyte population, as observed in the case of *Vav-Itgav*^*−/−*^ spleen tissues (Fig. 6H,I). In confirmation of this observation, flow cytometry analysis showed a significant increase in the proportion of CD41_+_ cells in the spleen MNC population (Suppl. Fig. 6C).

**Figure 6.**
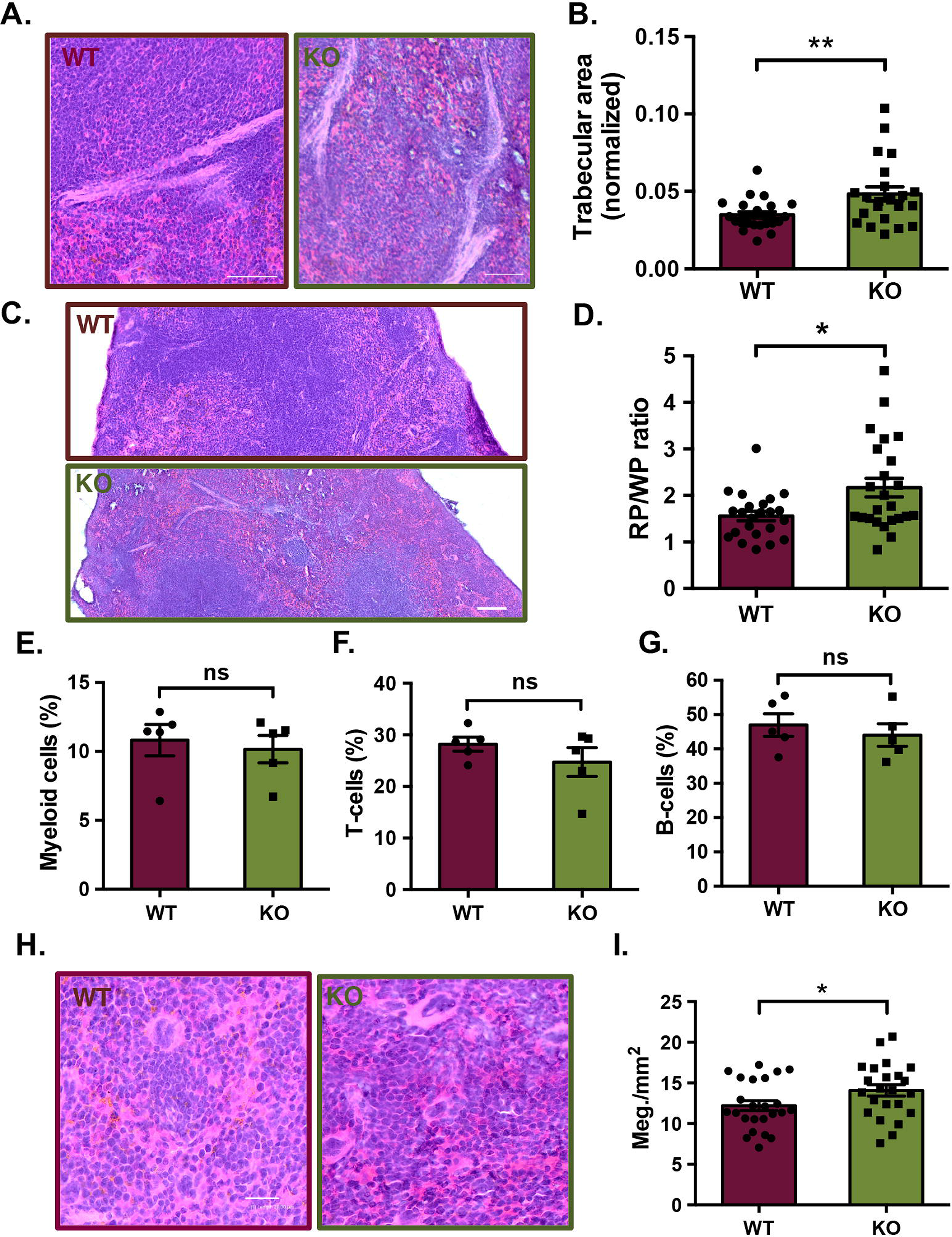
*Postn* deletion leads decline in spleen lympho-hematopoietic function. (A-D) Histological examination of spleen tissue sections from *Postn*^*+/+*^ (WT) and *Postn*^*−/−*^ (KO) mice. Tissues fixed in 4% paraformaldehyde were cut into 10**μ**m sections and H&E staining was performed. Brightfield images were captured for histological analysis. (A) Examination of the trabecular area by H&E staining. (B) Quantification of trabecular area as proportion of the total cross-sectional area of spleen tissues from WT and KO mice. (C) H&E stained sections analyzed for spleen cross-sectional area under red and white pulp. (D) Comparison of the ratio of red versus while pulp in the spleen tissues from WT and KO mice. n=4, N=23-24 (E-G) Flow cytometry analysis of the spleen mononuclear cells for the proportion of (E) myeloid cells (CD11b^+^/Gr-1^+^), (F) T cells (CD4^+^/CD8^+^), (G) B-cells (B220^+^). (H) Histological examination of the WT and KO spleen sections to identify megakaryocytes on the basis of morphological features. (I) Comparison of megakaryocyte frequency in spleen sections per mm2 of the total cross-sectional area. n=5, unpaired two tailed Student’s t-test was performed, * p<0.05, ** p≤0.01, ns indicates not significant.

### POSTN deficient spleen provides poor support to transplanted HSCs

Integrins are known for their role in cellular attachment in several cell types, including the hematopoietic system. Our studies published before, did not find any difference in the adhesion of *Itgav* deficient HSCs in vitro, also reflected in their homing potential in vivo (12, 27). Therefore, we next examined if this association played any role in supporting the incoming HSCs following BM transplantation in lethally irradiated mice. We performed colony forming unit-spleen assay (CFU-S_12_), wherein we transplanted healthy BM derived cells into WT (*Postn*^*+/+*^) or KO (*Postn*^*−/−*^) mice and compared the number of spleen colonies formed after 12 days (Fig. 7A,B). We normalised the number of colonies formed with the weight of the spleens and observed a significantly decreased number of spleen colonies in KO mice after 12 days of transplantation (Fig. 7C). We also compared the frequency of HSC sub-populations in the spleen tissues in the mice by flow cytometry analysis (Fig. 7D-G). While we observed a significant decrease in the frequency of lin-c-kit+ hematopoietic progenitors (Fig. 7E) and LSK cells (Fig. 7F), there was no difference in the frequency of the primitive HSCs (Fig. 7G). We did not observe any difference in the frequency of the lin-c-kit+ (Suppl. Fig. 5A), LSK cells (Suppl. Fig. 7B) and the primitive HSC populations (Suppl. Fig. 7C) of the BM. These results indicate that POSTN deficient spleen microenvironment might be poorer in its potential to maintain incoming hematopoietic progenitors.

**Figure 7.**
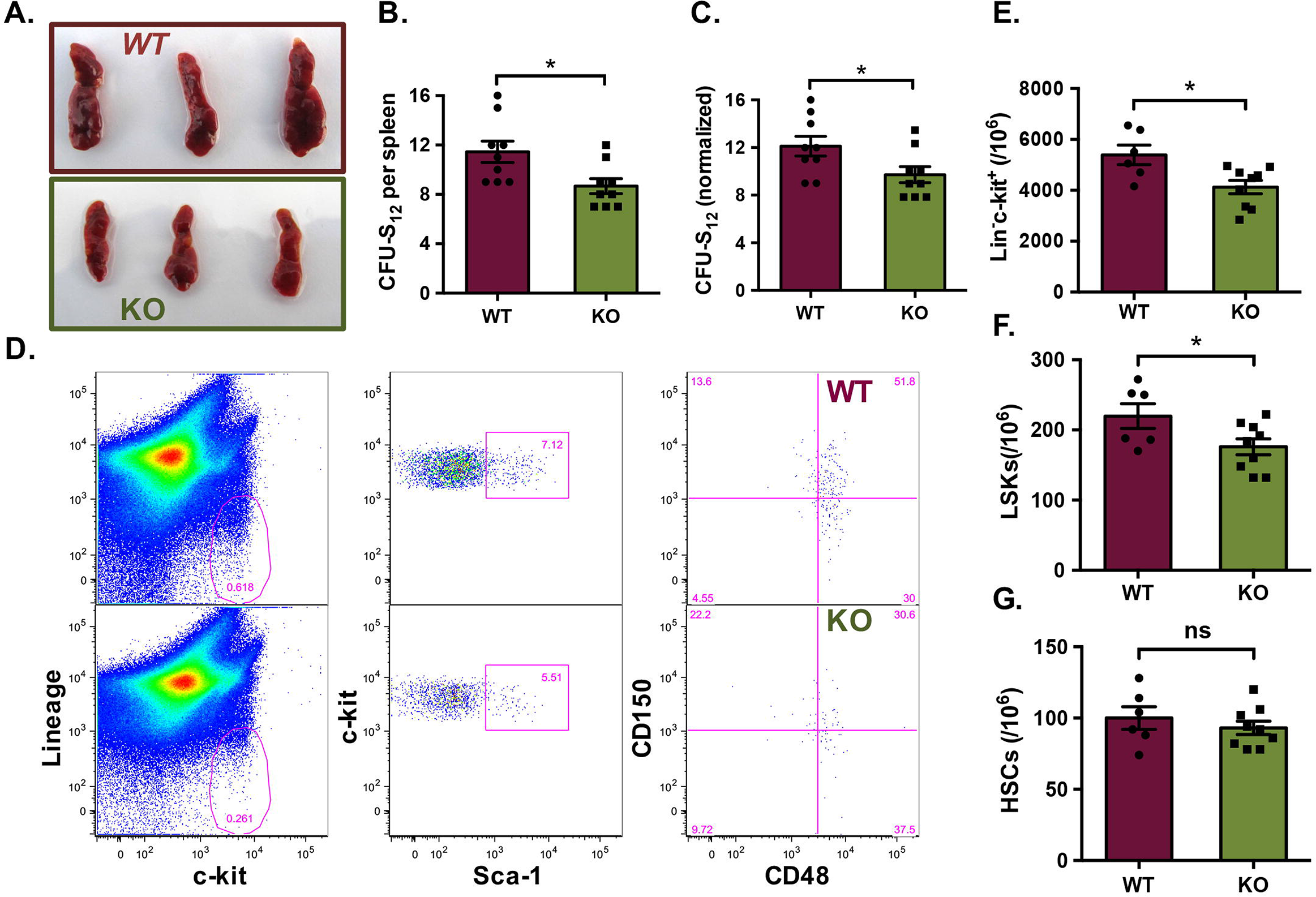
*Postn* deficiency leads to poor engra`ment/maintenance of HSCs. (A) CFU-S_12_ assay was performed to assess the HSC support potential of spleen. Overall morphology of the *Postn*^*+/+*^ (WT) and *Postn*^*−/−*^ (KO) spleen tissues, 12 days the mice received WT BM cells following lethal dose of irradiation. (B) Comparison of the total number of spleen colonies in recipient WT and KO mice from CFU-S_12_ assay. (C) Comparison of the number of spleen colonies in WT and KO mice following normalisation with the spleen weight. (D) Flow cytometry analysis of spleen MNCs for comparison of the HSC sub-populations in spleen, following CFU-S_12_ assay. Quantification of frequency of various hematopoietic stem and progenitor cell populations in the spleen from WT and KO mice; (E) lin-c-kit+ cells, (F) LSK cells, (G) primitive HSCs. Unpaired two tailed Student’s t-test was performed. n=3, N=9, t test: * p<0.05, ns indicates not significant.

## Discussion

We here present evidence for a role of POSTN-ITGAV interaction in spleen lympho-hematopoietic activity. Interruption in this interaction in *Postn*^*−/−*^ as well as *Vav-Itgav*^*−/−*^ led to increased proliferation rates in spleen HSC population. The impact of altered inside-out integrin signalling has been reported to affect spleen hematopoiesis through mobilization of hematopoietic progenitors due to altered BM hematopoietic niche (30). These effects could also be linked to poor homing of HSCs during embryonic development (31). Although, yolk sac (YS) did show the presence of *Itgb1−/−* HSCs in the chimeric mice, they were not detected in embryonic blood or the hematopoietic sites including spleen. Later, Alexander Medvinsky’s group through a comprehensive study showed that ITGA4, a key partner in integrin heterodimers with **β**1 chain, was not crucial for differentiation program in definitive HSCs (32). Furthermore, Talin1 that mediates association between integrin chains to actin cytoskeleton for cellular adhesion, was also shown to be dispensable for follicular B-cell maturation in spleen (33). While these and several other studies explored the role of inside-out integrin signaling in hematopoiesis, a role of outside-in integrin signalling in spleen hematopoietic function has not been characterised yet. In corroboration of our previous studies, we observed increased rate of HSC proliferation in both *Postn*^*−/−*^ and *Vav-Itgav*^*−/−*^ spleen. Importantly, methyl cellulose based colony assays showed a clear decline in their multi-lineage differentiation potential as fewer CFU-GEMMs were observed in *Vav-Itgav*^*−/−*^ spleen, despite an increase in the frequency of LSK and CD150+CD48-LSK cells.

It has been previously shown through in vitro experiments that POSTN plays a role in B-lymphopoiesis (34). Our previous study showed a decrease in the number of lymphocytes in PB of *Postn* deficient mice (12). The experiments presented in this study, clearly showed a decrease in the WP area indicating an effect on lymphopoietic activity. Flow cytometry experiments confirmed a B-cell specific effect upon *Vav-iCre* mediated loss of *Itgav*. Apart from an increase in the RP/WP ratio and a modest increase in overall spleen size, there was no indication of extramedullary hematopoiesis. Both *Postn*^*−/−*^ and *Vav-Itgav*^*−/−*^ mice showed increase in megakaryocyte numbers. In fact, there was a remarkable similarity in the phenotype of the two mouse lines used in this study. This, despite the fact that there are other known ligands for integrin heterodimers with ITGAV as one of the components. Several ECM proteins, soluble factors as well as ECM binding proteins such as Osteopontin (OPN) have been shown to bind heterodimers containing ITGAV (35). OPN is an established negative regulator of HSC function and *Opn−/−* mice showed increased numbers of HSCs in the BM as well as spleen (16). Under pathological conditions, soluble isoform of OPN increased lymphoid cell population in BM chimeras (36). In our earlier published work, *Vav-iCre* mediated deletion of *Itgav* led to decline in HSC function with increased proliferation rates. Our recent studies on fetal liver hematopoiesis also showed increase in proliferation rates both in the case of *Postn*^*−/−*^ and *Vav-iCre* embryos (27). Through the results presented in this manuscript, we further establish the role of POSTN-ITGAV interaction in the regulation of HSC proliferation.

We made another curious observation on spleen tissues from *Postn* deficient mice. The tissue showed an increase in the trabecular area. We did not observe this phenotype in *Vav-Itgav*^*−/−*^ spleen and hypothesised the presence of an additional hematopoietic-extrinsic effect of loss of *Postn*. This was corroborated by an overall small size of *Postn* deficient spleen, unlike in the case of *Vav-iCre* mediated *Itgav* deletion. Further proof came from CFU-S_12_ assays, wherein transplanted WT BM cells were poorly supported by the *Postn* deficient spleen niche and resulted in fewer spleen colonies. It is interesting to note that we observed POSTN expression mainly in the myofibroblastic cells of spleen trabeculae. Increase in the trabecular area points towards a feedback regulatory mechanism connecting POSTN expression and function. Poorer support for the incoming HSCs observed through CFU-S_12_ assays could also have resulted due to altered physico-structural support from the spleen niche in *Postn*^*−/−*^ mice.

Niche regulation of spleen resident HSCs has not been worked out yet. Using extramedullary hematopoiesis model, Sean Morrison’s group through elegant experiments showed the involvement of *Tcf21*-expressing endothelial and stromal cells in supporting HSCs in spleen (37). HSCs were found to be located close to the *Tcf21*^*+*^ stromal cells in the red pulp of spleen. These cells expressed SCF and SDF-1**α**; conditional deletion of anyone of these two genes, resulted in severely reduced spleen EMH. Potential of spleen stroma to support hematopoietic activity through in vitro experiments had been described (38). These cells supported the lymphoid cells from BM, spleen, thymus and blood in secondary cultures and led to robust differentiation into dendritic cells. Spleen stromal cell lines were later established that showed potential to support the development of hematopoietic cells including immature dendritic-like cells (39). Recently, these stromal cells that showed a rather restricted support of hematopoietic processes were found equivalent to Sca-1^+^gp38^+^Thy1.2^+^CD29^+^CD51^+^ fraction of spleen stroma (40). However, mechanisms that regulate the function of spleen resident HSCs have not been uncovered thus far. Associated phenotype of increased trabecular area in spleen from *Postn*^*−/−*^ mice indicates a possible regulation by the physical components of the niche.

As there was no induction of EMH upon Postn-deletion, the phenotypes observed are considered to be mediated via effects on spleen resident HSCs. Although their physiological relevance in the regular supply of blood cells or even EMH is not clearly understood, at clonal level their in vivo function was found to be equivalent to that of the BM HSCs (20). In fact, it has been reported that suppression of BM hematopoiesis is not crucial for the induction of EMH and extrinsic factors could stimulate increase in the number of hematopoietic progenitors in spleen and liver (41). A double transgenic mouse co-expressing IL-6 and soluble IL-6R showed elevated levels of EMH in spleen in addition to liver, without impacting BM hematopoiesis. In contrast, *hck−/−Src−/−* mice showed activation of EMH followed by severely affected BM hematopoiesis due to extreme form of osteopetrosis (42). Therefore, it has been challenging to understand the regulatory mechanisms that are played specifically in the spleen and their physiological relevance. Our report shows the involvement of outside-in integrin signaling not only in the regulation of spleen HSC function but possibly also in the creation of hematopoietic niche in this tissue.

## ACKNOWLEDGEMENT

This work was supported by the Wellcome Trust/DBT India Alliance Fellowship (IA/I/15/2/502061) awarded to SK and intramural funds from Indian Institute of Science Education and Research Thiruvananthapuram (IISER TVM). Institutional animal facility is supported by funds from the Department of Science and Technology, Government of India (under FIST scheme; SR/FST/LS-II/ 2018/217). SHM and AB are supported by IISER TVM. IMR is supported by Senior Research Fellowship from University Grants Commission (UGC), India. DP received support from INSPIRE fellowship from DST, India. CMV is supported by funds from KU Leuven (IDO/13/016 HSC-Niche) and FWO (G0E0117N).

## AUTHOR CONTRIBUTION

SK designed and supervised the study, wrote the manuscript. CMV provided material and reviewed the manuscript. SHM, IMR, AB and DP performed the experiments and analyzed the results. SS provided assistance with flow cytometry experiments. JH provided material.

