## Supplementary figures and images for "Niche mediated integrin signaling supports steady state hematopoiesis in spleen"

### Suppl. Fig. 1

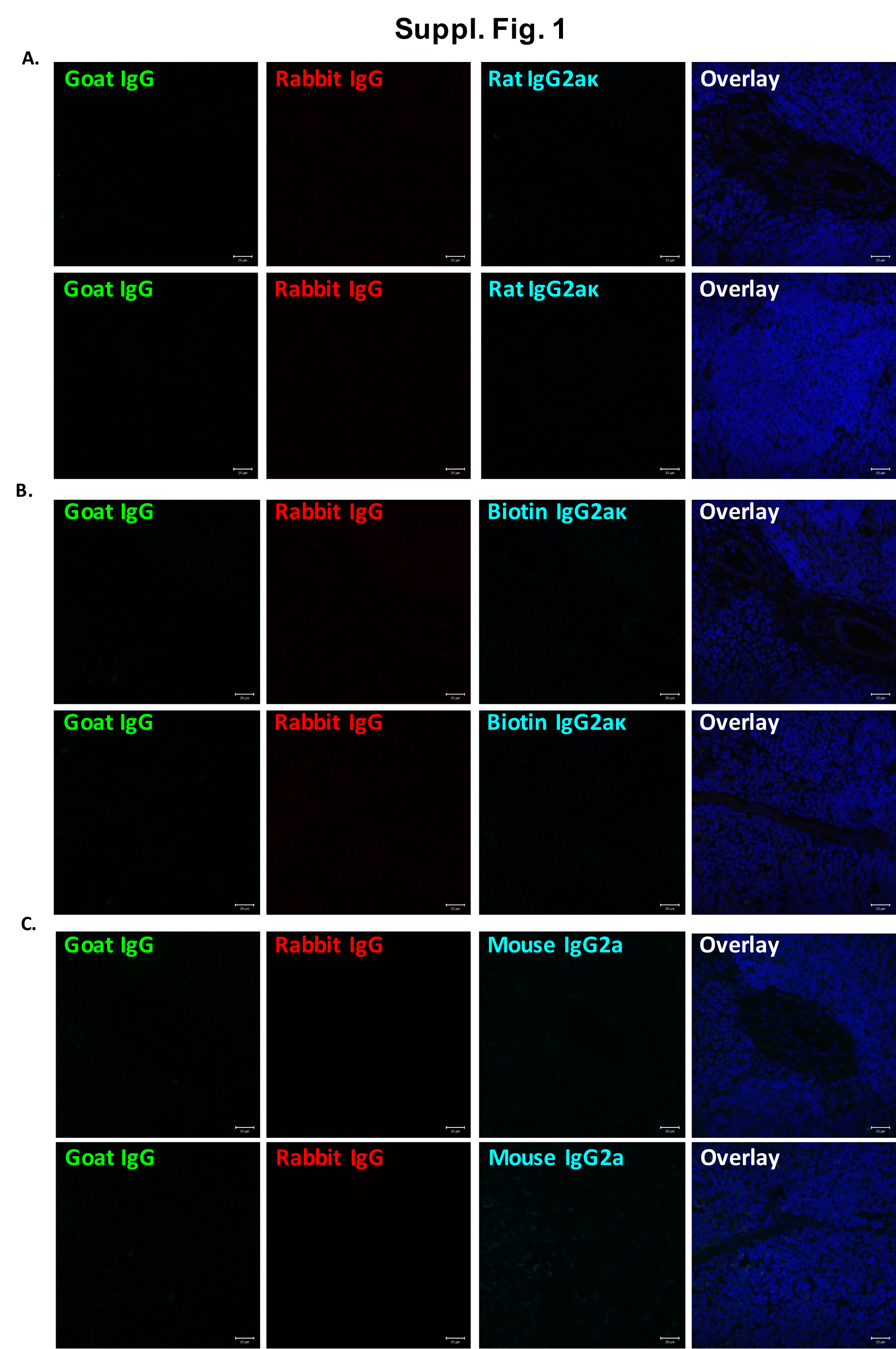

### Suppl. Fig. 2

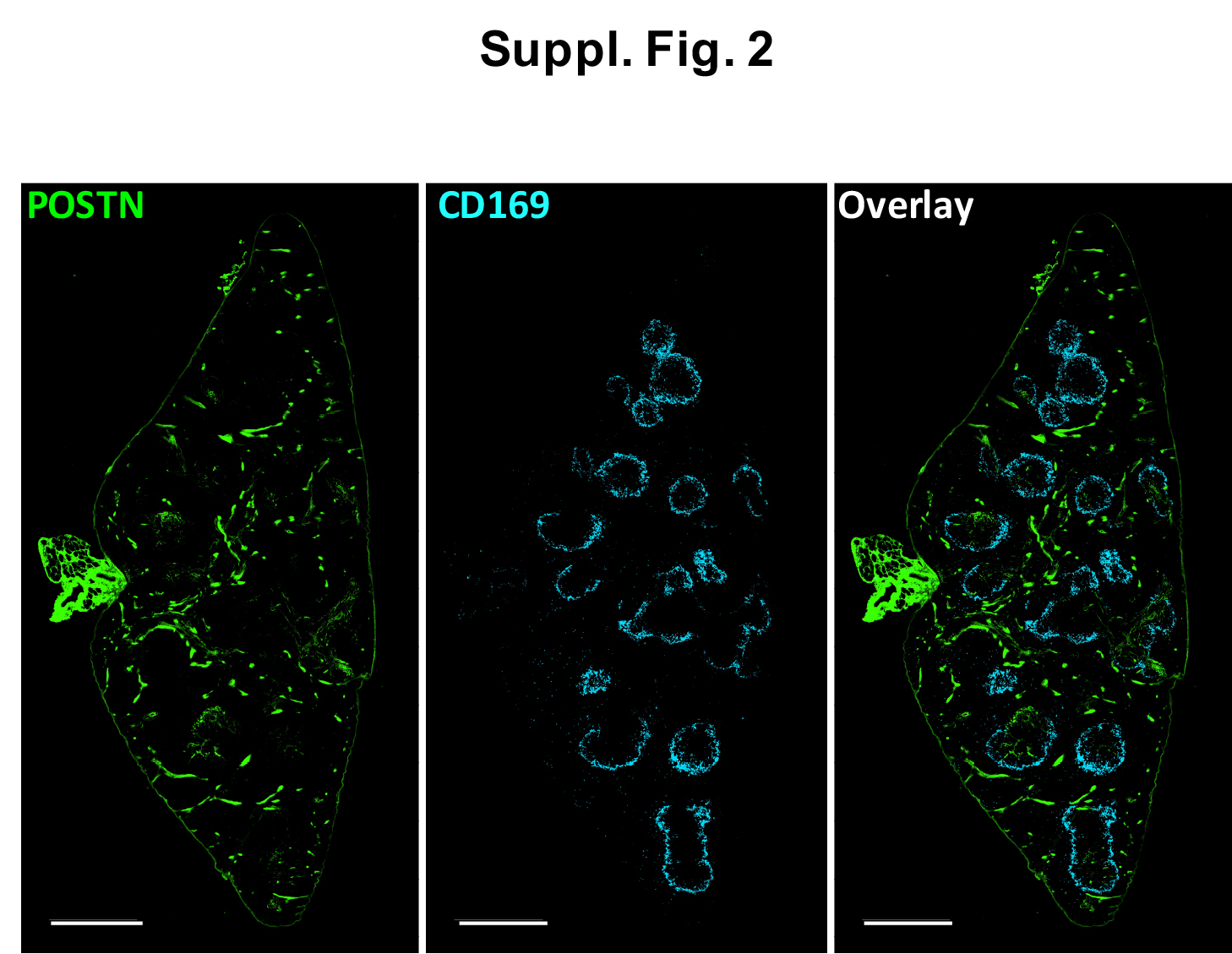

### Suppl. Fig. 3

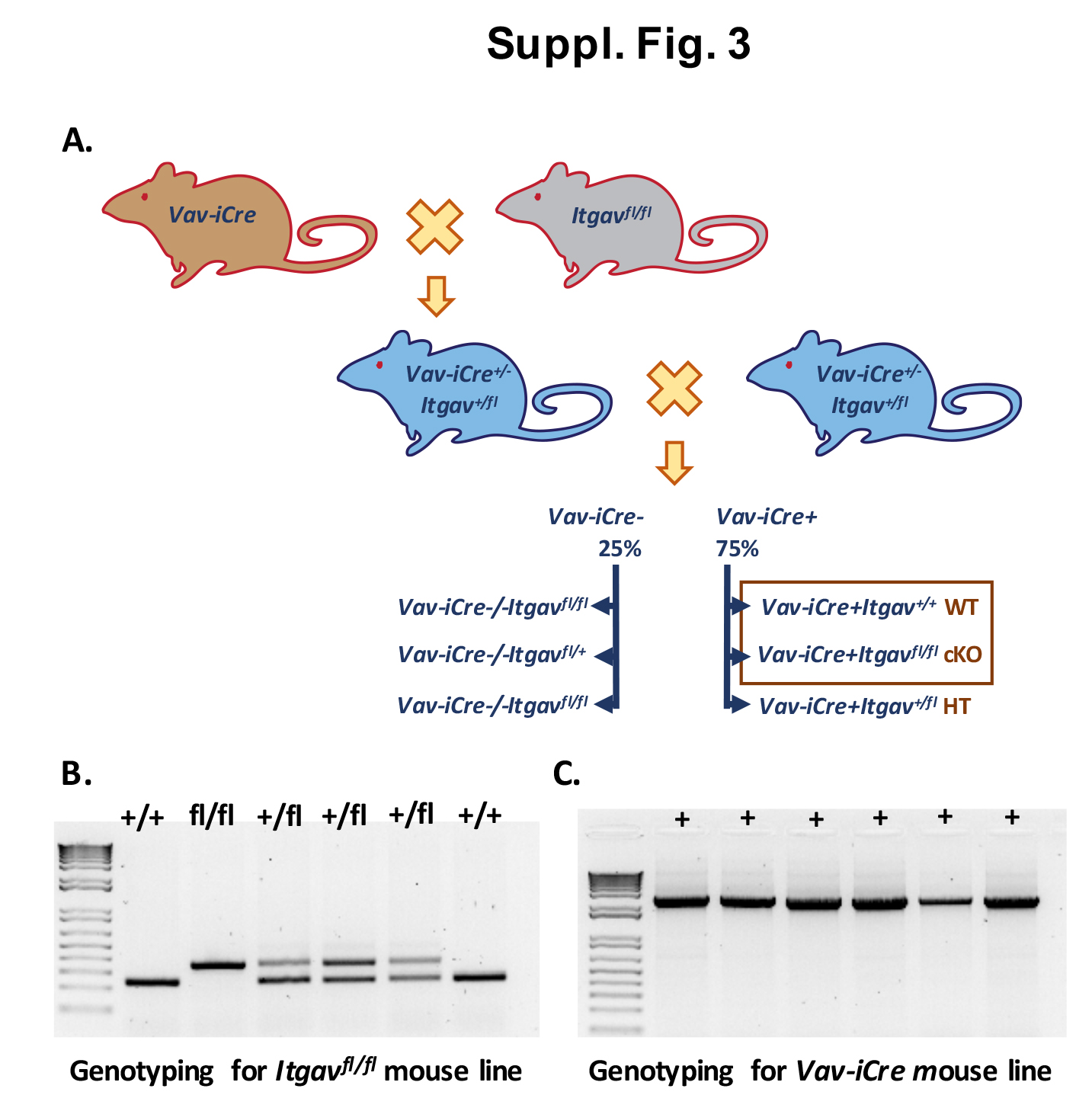

### Suppl. Fig. 4

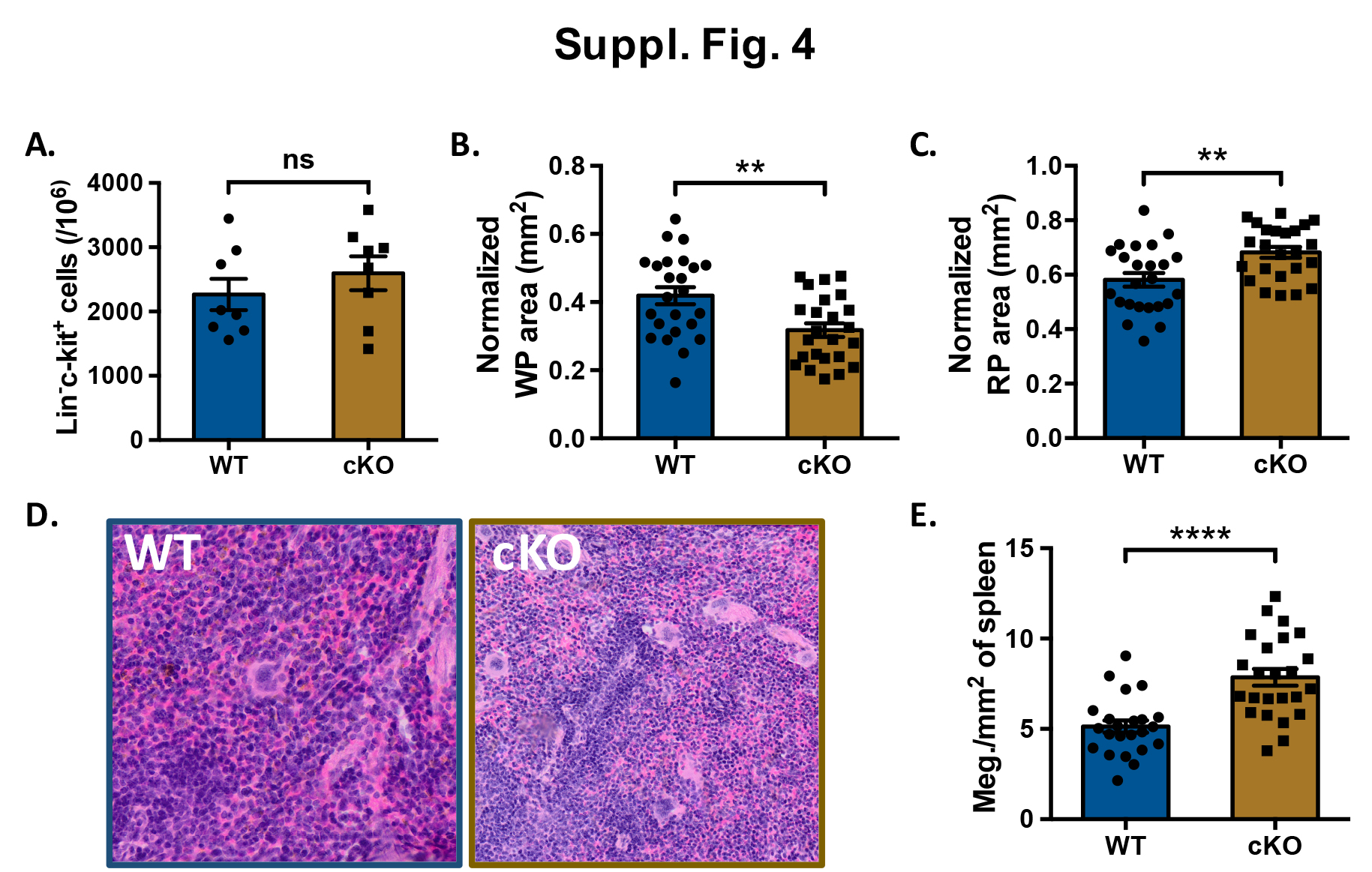

### Suppl. Fig. 5

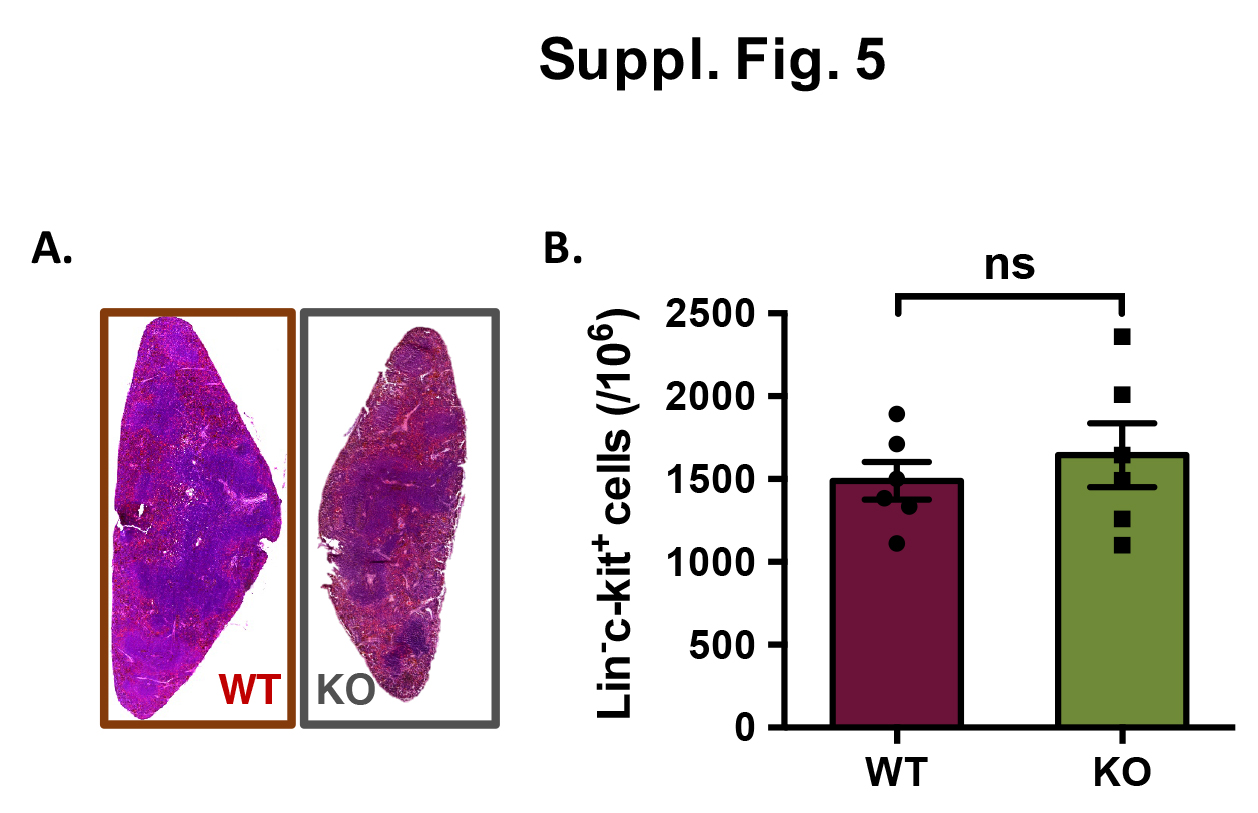

### Suppl. Fig. 6

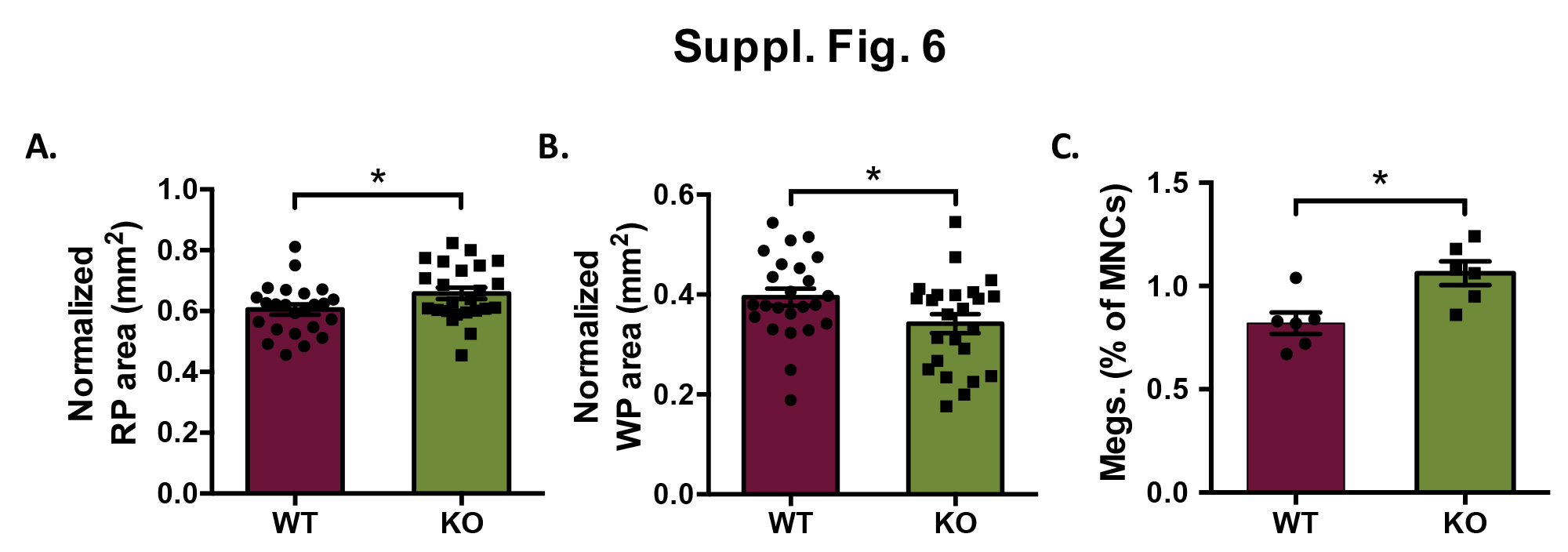

### Suppl. Fig. 7

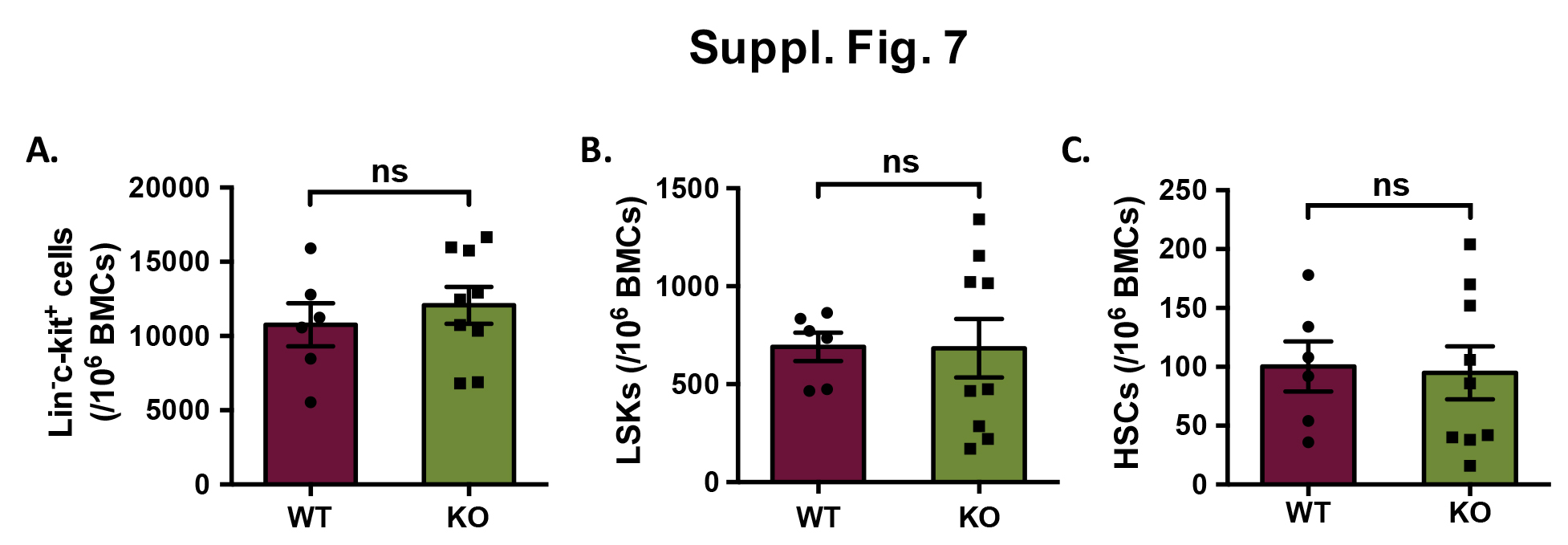
