## Supplemental material for "Niche mediated integrin signaling supports steady state hematopoiesis in spleen"

**Supplementary methods**

**Mice:** All procedures involving the use of animals were approved by the Institutional Ethics Committees (IAEC) of IISER Thiruvananthapuram and KU Leuven. At IISER Thiruvananthapuram, experiments were conducted following the guidelines of the Committee for the Purpose of Control and Supervision of Experimental on Animals (CPCSEA), government of India. *Itgav*$^{flox/flox}$ mice (Lacy-Hulbert et al., 2007) were crossed to *Vav-iCre* mice (from Thomas Graf, Centre for Genomic Regulation, Barcelona) to obtain *Vav-iCre;Itgav*$^{fl/fl}$ mice. Genotyping, as described before, was performed on genomic DNA from tail tip tissue. *Vav-iCre*$^{-}$*; Itgav*$^{fl/fl}$ and *Vav-iCre*$^{+}$*; Itgav*$^{+/+}$ littermates were used as controls. Sixteen-week old *Postn*$^{+/+}$ and *Postn*$^{-/-}$ mice, maintained, bred and genotyped as described before (Malanchi et al., 2012), were used in this study. The complete list of primers used for genotyping is provided in supplemental table 1. During the experiments, mice were maintained in isolator cages, fed with autoclaved acidified water and irradiated food ad libitum.

**Flow cytometry:** Hematopoietic stem and progenitor cell (HSPC) populations and lineage analysis of spleen derived cells was performed by flow cytometry. Lineage specific antibodies used were FITC conjugated Mac-1 and Gr-1 for myeloid cells, PECy7 conjugated B220 for B-cells, PE conjugated anti-CD4 and anti-CD8 for T-cells were used. All antibodies were procured from BD Pharmingen at were used at 0.25µg/ml concentration. Flow cytometric analysis for HSPC populations was performed using APC conjugated lineage antibody cocktail, PE conjugated anti mouse c-kit, PerCPCy5.5 conjugated anti-mouse Sca-1, FITC conjugated anti-mouse CD48 and PECy7 conjugated CD150 (0.25µg/ml; ebiosciences) antibodies. ITGAV and ITGB3 expression in the spleen derived HSCs was analyzed by using α-mITGAV and α-mITGB3 antibodies along with primitive HSC markers. Suitable isotype controls for each antibody were used in all experiments. The cells were analyzed by flow cytometry using FACS Aria III (BD Biosciences, San Jose, CA).

**Cell Cycle analysis:** BM derived cells were first stained for HSPC markers (Lineage, Sca-1 and c-kit) followed by Hoechst 33342 (Ho) in combination with Pyronin Y (PY) staining as described before (Arai et al., 2004). The cells stained for cell surface markers were incubated with Ho (10µg/ml) at 37°C for 45 min, followed by incubation with 1µg/ml PY for 30 min. Cells were acquired on FACS Aria III (BD Biosciences) and analysed using FlowJo software (TreeStar, Ashland, OR).

**Colony-forming unit-spleen assays:** Recipient *Postn*$^{+/+}$ and *Postn*$^{-/-}$ mice were lethally irradiated (10Gy) and injected with $1x10^5$ control FVB/NJ mouse BM derived cells. Three recipient animals were used for each donor mouse and the experiment was repeated thrice. Recipients were sacrificed 12 days after injection. Spleen colonies were visualized and counted as described before (Till and Mc, 1961).

**Histopathology:** The spleen tissues were harvested, washed in PBS and fixed with 4% paraformaldehyde (PFA) overnight at 4°C and washed with PBS and stored. The fixed tissues were dehydrated through gradient of ethanol and were cleared using xylene followed by paraffin infiltration. Paraffin blocks were made by embedding tissue in molten Paraplast X-Tra® using a Leica EG1150 Modular Tissue Embedding Centre. For histological analysis, 10µm thick tissue sections were cut using a microtome (Leica RM2265). FFPE sections on glass slides were stained with Harris' hematoxylin (HHS16; Sigma Aldrich) and eosin solution (HT110116; Sigma Aldrich). The sections were mounted using DPX mouting medium (DB6DF66104; MERCK) for microscopy. Brightfield images were captured on an Olympus IX83 inverted microscope, using 40X objective (UPlan FL N_40x/0.75_∞/0.17/FN26.5). CellSens Dimension 2.3 (Build 18987) software was used to capture and analysis of images.

**Immunohistochemistry and imaging:** Spleen tissues were harvested and washed in PBS and fixed with 4% paraformaldehyde (PFA) for 2 hours on shaker at ice-cold temperature and washed with PBS three times at 10 minutes interval and stored in 0.01% Sodium azide (Sigma_71289-50G) solution in PBS at 4°C. Fixed tissues were subjected to 48 hours 30% sucrose gradient for cryo-protection prior to cryo-block preparation using Polyfreeze (Sigma_P0091). Cryotome (Thermo Scientific HM525 NX) was used to obtain 10µm thick sections on frosted slides (VWR_631-0108). Immuno-labelling was performed using antigen specific primary antibodies; anti-periostin antibody (R&D systems, Minneapolis, MN), anti-α-smooth muscle actin antibody (abcam), anti-CD31 antibody (R&D systems, Minneapolis, MN), anti-collagenIV antibody (abcam), anti-laminin antibody (abcam) and fluorophore-conjugated secondary antibodies (Jackson ImmunoResearch Laboratories Inc.) Hoechst 33342 (Sigma) was used to counterstain the nuclei. Sections were mounted using Prolong Gold antifade mounting medium (Invitrogen_P36934). Fluorescence imaging was performed using Leica TCS SP5 II upright confocal microscope. Images were captured using 63X oil immersion objective (HCX PL APO CS 63.0x1.40 Oil) by software LAS AF. Leica Application Suit X (Version: 3.3.016799) software was used for analyzing the obtained images.

**Statistical analysis:** All data are represented as mean ± s.e.m. Comparisons between samples from two groups were done using unpaired Student's *t* test. For multiple comparisons, one-way ANOVA followed by Tukey Kramer post hoc test was used. Statistical analyses were performed with Microsoft Excel or GraphPad Prism 6. For all analyses, p-values <0.05 were accepted as statistically significant.

### Figure legends

**Supplementary Figure 1. Isotype controls for Immunostaining based experiments**

A) Isotype control for anti-mouse POSTN (Goat IgG), anti-mouse α-SMA (Rabbit IgG) and anti-mouse CD169 (Rat IgG2aκ) antibodies. Upper panel shows vascular region and lower panel shows white pulp region corresponding to the ones presented in Figure 1A-D.

(B) Isotype control for anti-mouse POSTN (Goat IgG), anti-mouse α-SMA/Laminin/Col IV (Rabbit IgG) and anti-mouse CD31 (Biotin IgG2aκ). Upper panel shows vascular region and lower panel shows trabecular region corresponding to the ones presented in Figure 1E and 2A.

(C) Isotype control for anti-mouse POSTN (Goat IgG), anti-mouse Laminin/Col IV (Rabbit IgG) and anti-mouse α-SMA (Mouse IgG2a). Upper panel shows vascular region and lower panel shows trabecular region corresponding to the ones presented in Figure 2B,C.

(scale bar=20μm).

**Supplementary Figure 2. POSTN is expressed in myofibroblasts in adult spleen**

POSTN expression was examined in the adult spleen tissues using immunohistochemistry on 10µm cryo-sections followed by fluorescence imaging using tile scanning. Specific antibodies were used to identify the cells expressing POSTN, macrophages lining the WP areas were identified by using CD169 staining, nuclear counterstaining was done using Hoechst 33342.

(n=2, N=8, scale bar=0.5mm).

**Supplemental Figure 3. Vav-iCre mediated deletion of Itgav**

(A) Schematic representation of breeding scheme used to generate *Vav-iCre*$^{+}$*;Itgav*$^{+/+}$ (WT) and *Vav-iCre*$^{+}$*;Itgav*$^{-/-}$ (cKO) mice. *Vav*-iCre and *Itgav*$^{fl/fl}$ mice were crossed to delete *Itgav* in the hematopoietic cells.

(B,C) Genotyping PCR performed on tail-tip DNA to identify *Vav-iCre*$^{+}$*;Itgav*$^{+/+}$ (WT), *Vav-iCre*$^{+}$*;Itgav*$^{fl/+}$ (HT) and *Vav-iCre*$^{+}$*;Itgav*$^{-/-}$ (cKO) mice. PCR was performed to identify *Vav-iCre* (B) and Itgav$^{fl}$/Itgav$^{+}$ (C) alleles, separately.

**Supplemental Figure 4. *Itgav* deletion leads to poorer lymphopoietic function**

(A) Mononuclear cells from *Vav-Itgav*$^{+/+}$ (WT), *Vav-Itgav*$^{-/-}$ (cKO) spleen tissues were used for the analysis of HSPC (lin$^{-}$c-kit$^{+}$ cells) frequency by flow cytometry (n=8).

(B,C) Spleen tissues isolated from WT and cKO animals were harvested. The formalin-fixed, paraffin-embedded tissues were used to cut 10μm sections that were used for H&E staining. (B) The total cross-sectional area under red pulp (RP), and (C) white pulp (WP) was compared between WT and cKO spleen tissues (n=4, N=24).

(D) Histological examination of the WT and cKO spleen sections to identify megakaryocytes on the basis of morphological features.

(E) Comparison of megakaryocyte frequency in spleen sections per $mm^2$ of the total cross-sectional area. (n=4, N=24)

Unpaired two tailed Student's t-test was performed. * $p<0.05$, ns indicates not significant.

**Supplemental Figure 5. *Postn* deletion leads to poorer lymphopoietic function**

(A) Comparison between the *Postn*$^{+/+}$ (WT) and *Postn*$^{-/-}$ (KO) mice for total spleen cross-sectional area. Spleen tissues were harvested, formalin-fixed and cut into 10μm sections used for H&E staining.

(B) Comparison between the WT and KO spleen tissues for the frequency of hematopoietic stem and progenitor cell population (lin$^-$c-kit$^+$ cells) by flowcytometry.

**Supplemental Figure 6. *Postn* deletion leads to poorer lymphopoietic function**

(A,B) Spleen tissues isolated from WT and KO animals were harvested. The formalin-fixed, paraffin-embedded tissues were used to cut 10μm sections that were used for H&E staining. (A) The total cross-sectional area under red pulp (RP), and (B) white pulp (WP) per $mm^2$ of the total spleen cross-sectional area was compared between WT and KO spleen tissues (n=4, N=24).

(D) Flow cytometry analysis of the spleen mononuclear cells (MNCs) for the frequency of megakaryocytes identified as CD41$^+$ cells using specific antibodies.

**Supplemental Figure 7. *Postn* deficiency doesn't affect short-term stem cell homing in the BM**

CFU-$S_{12}$ assay was performed to assess the HSC support potential of spleen. *Postn*$^{+/+}$ (WT) and *Postn*$^{-/-}$ (KO) mice received WT BM cells following lethal dose of irradiation. After 12 days of grafting, spleen colonies were enumerated along with BM analysis for the analysis of stem cell sub-populations. Flow cytometry analysis of spleen MNCs for comparison of the HSC sub-populations in BM. Quantification of frequency of various hematopoietic stem and progenitor cell populations; (A) lin$^-$c-kit$^+$ cells, (B) LSK cells, (C) primitive HSCs.

Unpaired two tailed Student's t-test was performed. n=3, N=9, t test: ns indicates not significant with $p \geq 0.05$.

### Table 1

| Sr. No. | Antibody | Clone | Isotype | Company | Catalog no. |
|---|---|---|---|---|---|
| For microscopy | | | | | |
| 1 | Anti-mouse Periostin | Polyclonal | Goat IgG | R&D | AF2955 |
| 2 | Anti-mouse α-smooth muscle actin | Polyclonal | Rabbit IgG | Abcam | ab5694 |
| 3 | Anti-mouse CD31 | Polyclonal | Goat IgG | R&D | AF3628 |
| 4 | Biotin anti-mouse CD31 | MEC13.3 | Rat IgG2a,κ | Biolegend | 102504 |
| 5 | Anti-Collagen-IV | Polyclonal | Rabbit IgG | abcam | ab6586 |
| 6 | Anti-Laminin | Polyclonal | Rabbit IgG | abcam | ab11575 |
| 7 | Anti-CD169 | 3D6.112 | Rat IgG2a,κ | Biolegend | 142419 |
| 8 | Anti-α-smooth muscle actin | 1A4 | Mouse IgG2a | R and D | MAB1420 |
| 9 | Fab Fragment | Polyclonal | Goat IgG | Jacksons. Imm Res. | 115-007-003 |
| For flowctometry | | | | | |
| 1 | Anti-mouse Gr-1 FITC | RB6-8C5 | Rat IgG2b, κ | eBioscience | 11-5931-85 |
| 2 | Anti-mouse CD3e FITC | 145-2C11 | Armenian Hamster IgG | eBioscience | 11-0031-85 |
| 3 | Anti-mouse B220 FITC | RA3-6B2 | Rat IgG2a, κ | eBioscience | 11-0452-82 |
| 4 | Anti-mouse Ter119 FITC | TER-119 | Rat IgG2b, κ | BD Biosciences | 557915 |
| 5 | Anti-mouse CD11b FITC | M1/70 | Rat IgG2b, κ | BD Biosciences | 557396 |
| 6 | Anti-mouse CD48 FITC | HM48-1 | Armenian Hamster IgG | eBioscience | 11-0481-85 |
| 7 | Anti-mouse CD48 APC-Cy7 | HM48-1 | Armenian Hamster IgG | BioLegand | 103432 |
| 8 | Anti-mouse c-kit APC | 2B8 | Rat IgG2b, κ | BioLegand | 105812 |
| 9 | Anti-mouse c-kit APC-Cy7 | 2B8 | Rat IgG2b, κ | BioLegand | 105826 |
| 10 | Anti-mouse CD51 PE | RMV-7 | Rat IgG1, κ | BD Biosciences | 551187 |
| 11 | Anti-mouse CD61 PE | 2C9.G3 | Armenian Hamster IgG | eBioscience | 12-0611-83 |
| 12 | Anti-mouse CD61 AF647 | 2C9.G2 (HMβ3-1) | Armenian Hamster IgG | BioLegand | 104314 |
| 13 | Anti-mouse Sca-1 BB700 | D7 | Rat IgG2a, κ | BD Biosciences | 742089 |
| 14 | Anti-CD150 PE-Cy7 | TC15-12F1 2.2 | Rat IgG1 | BioLegand | 115914 |

**Table 2**

| Primer name | Primer sequence (5’-3’) |
|---|---|
| Genotyping Postn mice | (F) GGT GCT TCT GTA AGG CCA TC<br>(R) GTG AGC CAG GAC CTT GTC ATA<br>(Int-as) AGC ACT GAC TGC GTT AGC AA |
| Genotyping Itgav mice | (F) GGTGACTCAATCTGTGACCTTCAGC<br>(R) CACAAATCAAGGATGACCAAACTGAG |
| Genotyping Vav-iCre mice | (F) CCATGGCACCCAAGAAGAAG<br>(R) GCTTAGTTTTCCTGCAGCGG |
